# Transient optical clearing using absorbing molecules for ex-vivo and in-vivo imaging

**DOI:** 10.1101/2025.04.02.646849

**Authors:** Muhammed Waqas Shabbir, Matthew Phillips, David Asante-Asare, Zihao Ou

**Affiliations:** Department of Physics, The University of Texas at Dallas, Richardson, TX 75080; Department of Electrical and Computer Engineering, The University of Texas at Dallas, Richardson, TX 75080

## Abstract

Dynamic imaging is a foundational tool in biology and medicine but is limited by light scattering in live tissues caused by refractive index mismatches, which reduces penetration depth and resolution in biological tissues. Our previous work introduced a novel approach to achieve an optical transparency in live animals by utilizing strongly absorbing molecules. When these absorbing molecules, such as Tartrazine, an FDA approved dye commonly used in foods, dissolve in water, they modify the refractive index of the aqueous medium through Kramers-Kronig relations. This modulation allows reduction of the refractive index mismatch between separate components in tissue with distinct refractive indices, thus mitigating scattering. In this article, we report detailed methodology of our tissue clearing technique using a Tartrazine-based aqueous solution, applied to both ex-vivo and in-vivo samples. This approach reversibly renders live mice bodies transparent in the visible spectrum, enabling clearer and deeper imaging of internal processes and structures. We then utilized an advanced image processing algorithm to improve the contrast and segmentation of organs of interests. We also show high transparency in ex-vivo samples, such as chicken breast, when treated with Tartrazine based solution. This method offers a versatile and accessible approach to enhancing imaging capabilities across various modalities and paves the way for advancement in our understanding of complex biological systems.

**SUMMARY:** Optical clearing techniques, such as absorbing molecules solutions, reduce light scattering temporarily to enhance biological imaging and offer biocompatible, reversible in-vivo imaging strategies for deeply embedded organs. This protocol presents detailed experimental procedures as well as advanced image processing to achieve high-quality, dynamic imaging for real-time dynamics in living animals.

## INTRODUCTION

Biological imaging has long been challenged by the scattering of light, which obscures structures of interest.^1^ Optical clearing techniques aim to reduce light scattering and enhance visual clarity, enabling the visualization of previously hidden structures.^2^ The development of these techniques has led to a diverse array of methods designed to improve optical imaging of ex-vivo biological tissues, even whole animals, without physical dissection of the samples.^3^ However, development of similar techniques for in-vivo applications has been challenging.

Techniques such as CLARITY offer high-resolution imaging but lack reversibility, making them unsuitable for live animal studies.^4^ Constantini et al. have shown slower progress of in-vivo imaging compared to ex-vivo imaging clarity, which is due to the complexity of maintaining tissue viability while avoiding toxic levels of agents needed for tissue transparency.^5^ Genina emphasizes the future of optical tissue clearing as reliant on non-invasive and reversible methods.^6^ Although ex-vivo tissue clearing methods like CLARITY and iDISCO achieve high optical transparency and subcellular resolution through lipid removal or solvent-based clearing, they are often unsuitable for in-vivo imaging with live animals.^4,7,8^ This limitation affects dynamic imaging, which is crucial for biomedical imaging of dynamic processes inside living animals, such as disease development and therapeutics.

Absorbing molecules appear promising for achieving optical transparency of tissues in living animals.^9^ By introducing strongly absorbing molecules like Tartrazine, a food-grade dye, the tissue increases absorption of light in the ultraviolet and blue portions of the spectrum, leading to an increased refractive index and reduced scattering for longer wavelengths. This counterintuitive optical tissue clearing technique is optimized for in-vivo imaging and the Kramers-Kronig relations provide a scientific foundation for the observed transparency when using strongly absorbing molecules like Tartrazine.^10,11^ For in-vivo imaging, Tartrazine is combined with heated agarose particles and water to create a cohesive solution that adheres to skin better. The water-soluble nature of Tartrazine also facilitates easy removal of the dye-based solution from live animals, reducing potential effects of extended exposure to research chemicals. Additionally, the method has been adapted for various biomedical applications to improve the imaging depth, resolution as well as sensitivity, including optical coherence tomography^12,13^ and photoacoustic imaging,^13,14^ demonstrating its potential for the broader biomedical imaging community.

With the growing interest in the application of absorbing molecules in biomedical imaging,^15,16^ the protocol presents a detailed procedure of utilizing food dye for ex-vivo and in-vivo applications, along with image processing and reversible in-vivo imaging. This protocol is applied to chicken breast tissue to demonstrate ex-vivo transparency and is then modified for in-vivo head and abdominal imaging of mice, showing reversibility. A solution applied to mice skin in-vivo creates temporary, reversible transparency without harming the skin or tissue integrity. This technique can be integrated with an image processing pipeline, including denoising, to create an optimized imaging protocol for in-vivo optical imaging of internal organs. When combined, these techniques enable better non-invasive, longitudinal imaging for real-time observations of biological processes. The use of safer agents that demonstrate reversibility overcomes previous limitations and provides a more suitable method for dynamic biological imaging.

## PROTOCOL

### 1. Tartrazine Solution Preparation

#### a) For ex-vivo samples

1. To prepare dye solutions, calculate the required mass of dye powder (Tartrazine) to prepare the desired volume of the solution. Use the formula. Mass (g) = Molarity (M) × Molecular Weight (g/mol) × Volume (L) Note: In this work, we use recalibrated molarity value. The recalibration is performed using the absorption spectra of dye solutions and Beer-Lambert law to quantify the molar concentrations.^17^ Caution: Use an analytical balance to weigh the precise amount of the powder.
2. Dissolve the dye in deionized water. Use volumetric pipettes for accuracy.
3. Heat solution at 40^0^C for 10 minutes if it is not completely dissolved in DI water at room temperature (**Figure 1a**). Low molarity solutions are completely soluble at room temperature, and remain in liquid form, while high concentration (> 0.37 M) solutions require heating. Caution: Excessive heating can evaporate water and alter the actual concentration of the solution.
4. Stir on vortex mixer for 30 seconds to one minute to ensure complete dissolution.
5. To prepare a 0.65 M solution, dissolve 427.488 mg tartrazine powder in 1 ml DI water.^17^ For other concentrations, see **Table 1**.

**Figure 1.**
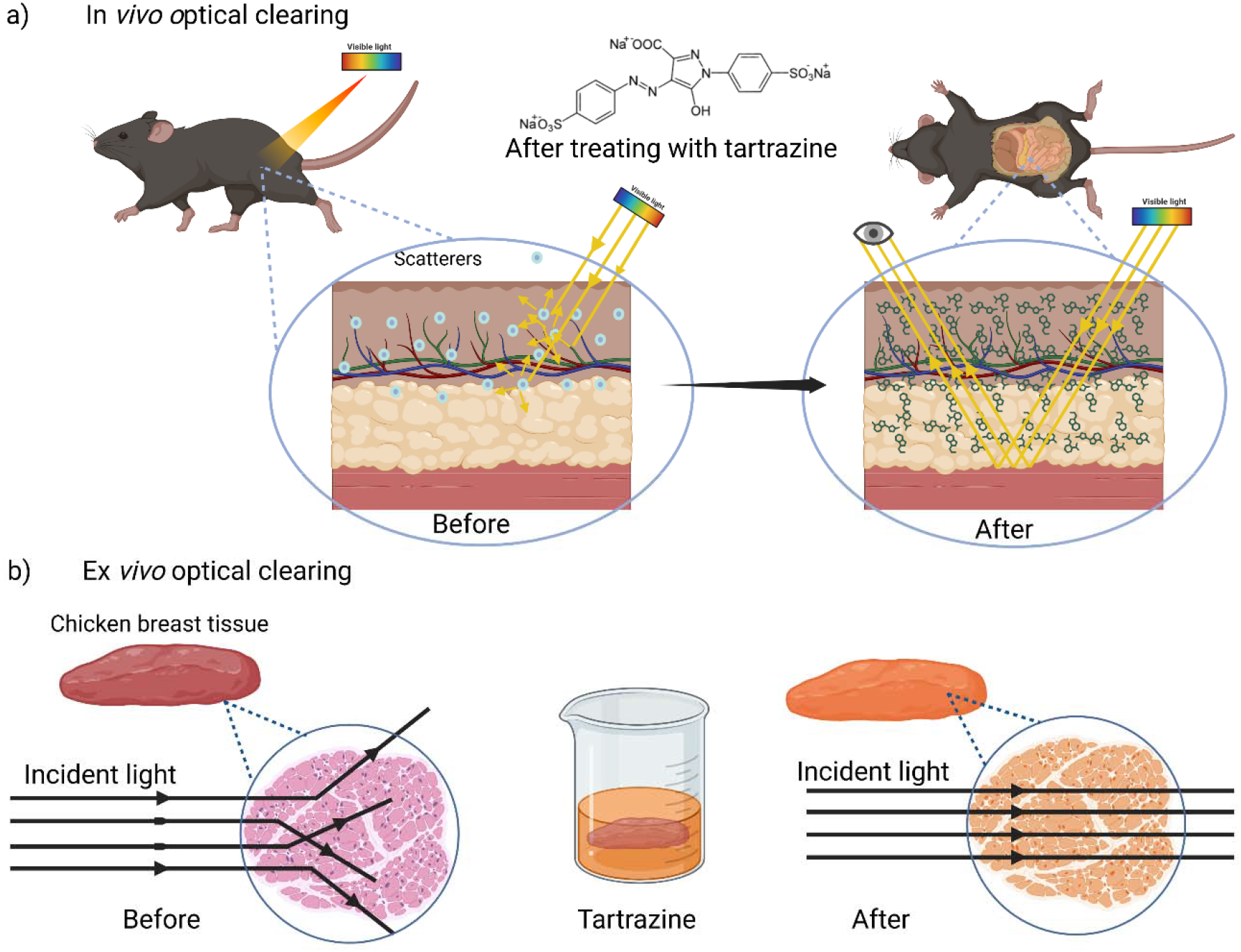
Schematic illustration of the in-vivo (**a**) and ex-vivo (**b**) optical clearing of tissue using absorbing molecule solutions.

**Table 1.** Preparation recipe of Tartrazine solutions of different concentrations.

| Tartrazine mass (mg)<br>dissolved in 1 mL water | Relative percentage of<br>tartrazine in solution | Calibrated molecular<br>concentration (M) |
| --- | --- | --- |
| 106.9 | 9.7% | 0.18 |
| 213.7 | 17.6% | 0.37 |
| 320.8 | 24.3% | 0.53 |
| 426.6 | 29.9% | 0.65 |

#### b) For In-vivo Samples

1. Calculate the required mass of low-melting temperature agarose microparticles and tartrazine to prepare a 0.65M concentration with 0.4% (4 mg/ml) agarose hydrogel solution.
2. Add deionized water and heat the solution at 75^0^C for approximately 10 minutes till the agarose particles are completely dissolved.
3. Occasionally mix using a vortex mixer during the heating process for uniform dissolution and allow the solution to cool down before use.
4. If the solution solidifies into gel form, use the vortex mixer for 30 seconds to re-liquefy it.
5. To prepare a 0.65M hydrogel solution, dissolve 427.488 mg Tartrazine powder and 4 mg agarose in 1 ml DI water.

### 2. Optical Clearing of Ex-vivo tissues

#### a) Sample Preparation

1. To prepare ex-vivo tissue, slice a chicken breast to the desired thickness parallel to collagen fibers.
2. Clean the sliced tissue with water and measure the thickness.
3. Soak the tissue in the dye solution.
4. Caution: Make sure that the volume of the solution is more than the sample volume and tissue is completely submerged.
5. Shake the sample for 10 minutes on an orbital shaker at 100 revolutions per minute.
6. Let the tissue stay in the solution for 60 minutes at room temperature for low concentrations ≤ 0.4M and at 40^0^C for higher concentrations > 0.4M.

#### b) Imaging

1. After removing the sample from the solution, sandwich it between two transparent glass slides and capture the images.
2. Conduct the imaging process in controlled light environment to minimize external light interference.
3. Repeat the imaging process for tissues with different thicknesses and solutions of different concentrations (**Figure 2b**). Note: In this work, we utilize LED light source for back illumination, but it works in the exact same way for top illumination light source as well).

**Figure 2.**
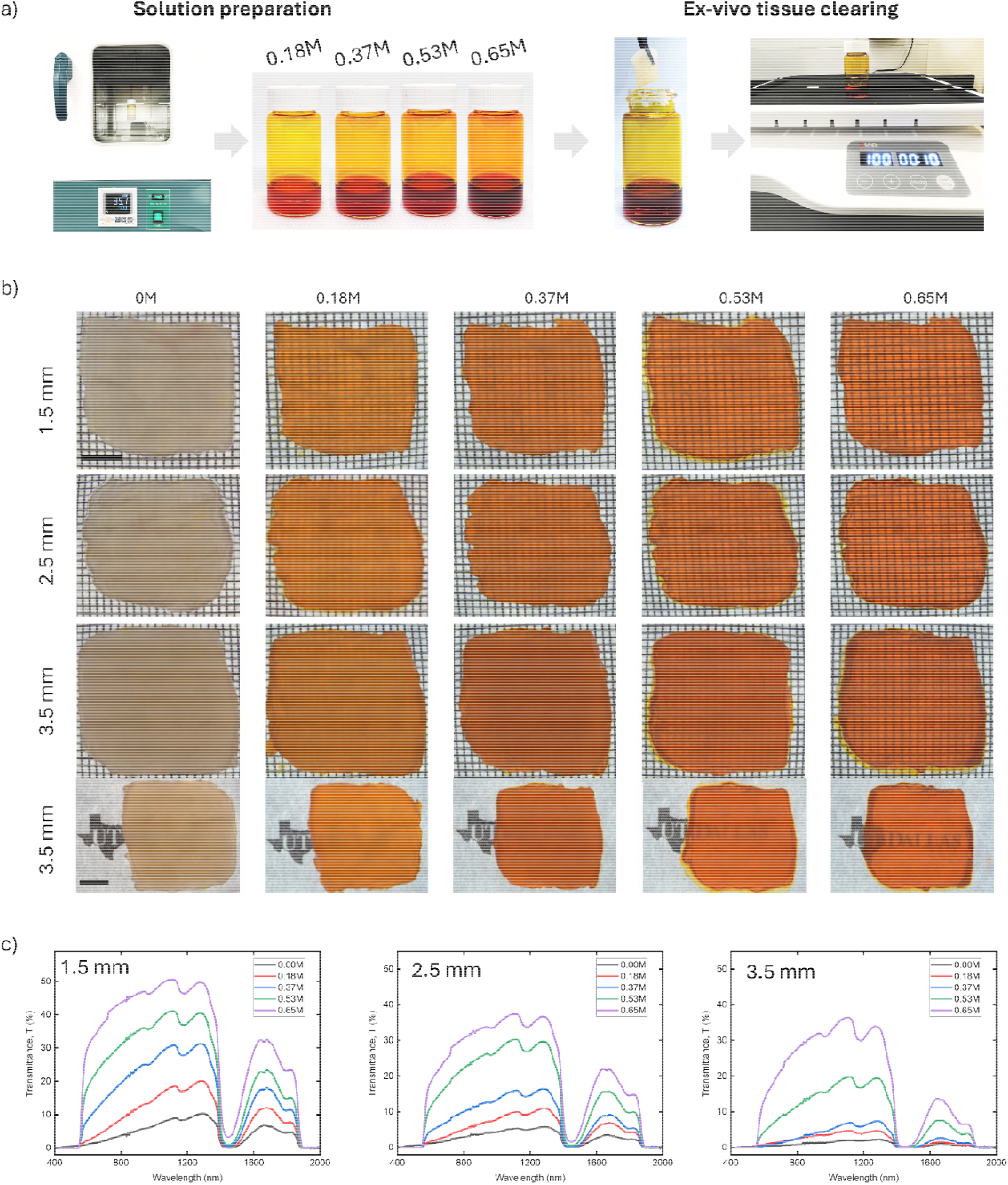
Protocols for optical clearing of ex-vivo tissue. (**a**) Photographs of the solutions preparation and clearing processes of the ex-vivo tissue. (**b**) Optical images of the chicken breast tissue with different thicknesses placed on top of a pattern. (**c**) Optical transmittance spectra of the chicken breast tissue with different thicknesses. All Scale bars: 5 mm.

#### c) Optical Transmission Measurements

1. To measure the optical transmission, sandwich the chicken tissue between two transparent glass slides and measure the transmission using UV-Visible-NIR Spectrometer (**Figure 2c**).
2. Make sure the system is baselined properly, and the light beam passes through the same region of the tissue before and after treating it with dye solutions. Note: The transmission spectra reported in this paper is measured across a 250nm to 2600nm wavelength range, using a medium scan speed and a data interval of 1.0 nm.

### 3 Optical transparency in live animals

#### a) Animal Preparation

1. Before the experiment, measure the weight of the animal and note down the activity level.
2. Use a heating pad with temperature set to 37^0^C to maintain the animal’s body temperature during the experiment.
3. Anesthesia is necessary for optical imaging of living animals. For abdominal imaging, inhalation anesthetics are recommended for quick recovery. For inhalation anesthesia, induce anesthesia using isoflurane at 1.5-3% and flow rate 0.8L/minutes for approximated 5 minutes before the imaging experiment and maintain it at 1-2% throughout the experiment. For injectable anesthesia, prepare a cocktail solution of containing Ketamine HCL (16 mg/mL) and Dexmedetomidine HCL 0.2mg/mL.
4. Trim the hair from the belly for gut imaging and from the head for brain imaging.
5. Apply depilatory cream (Nair), and make sure that Nair is not applied to the skin for more than 60 seconds to avoid irritation.
6. Clean the skin with sterile alcohol wipes.
7. Use Atipamezole as a reversal drug for injectable anesthesia after the experiment.

#### b) Procedure for making Tissue Transparent

1. Prepare the imaging set up, heating pad set to 37^0^C to avoid hypothermia, aligning the light source, and camera to capture the images.
2. Use a 0.65 M Tartrazine solution mixed with 0.3-0.4% agarose gel for application.
3. Apply the solution to the skin (belly or head, depending on the nature of the experiment) using a dropper and allow it to sit for a few seconds for absorption (**Figure 3a**).
4. Gently scrub the area using a Q-Tip for even distribution.
5. Massage the area gently with a Q-Tip and apply more solution till the skin starts showing transparency and internal organs are visible. Caution: Ensure that the solution does not crystallize over the application area, crystallization can block skin passages hindering the dye molecules absorption in the skin, which is highly crucial to achieve transparency. Sufficient absorption of dye molecules turns skin’s color to orange red.
6. Continue the process of massaging until the skin starts showing transparency and internal organs are visible.
7. Capture the images using a camera ensuring proper focus and exposure.
8. Place a glass slide or cover slip on the skin to minimize shine and reduce reflection for better quality images.

**Figure 3.**
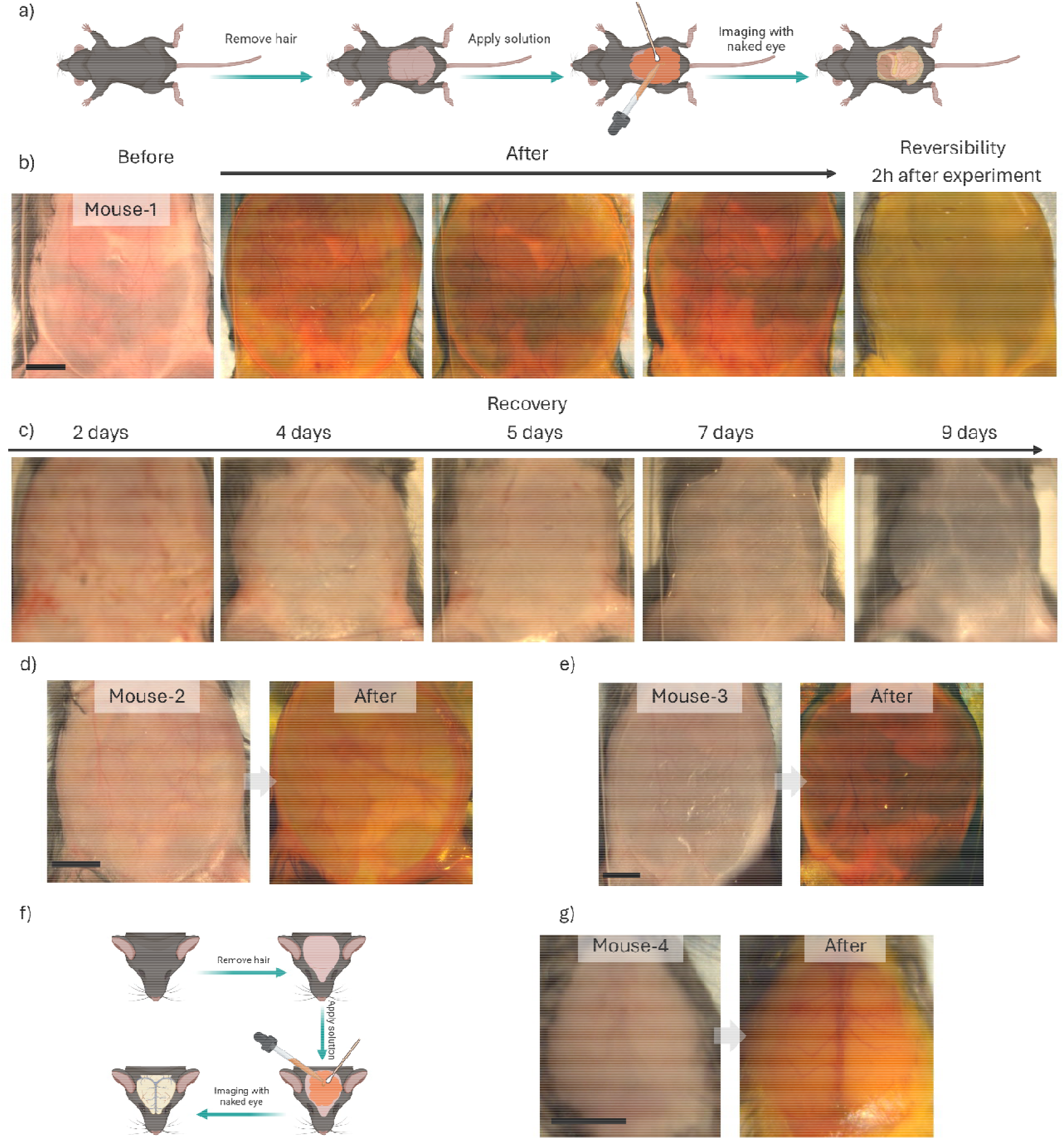
Protocols for optical clearing and imaging of animals. (**a**) Schematics illustrating the application of absorbing molecules on the mouse belly. (**b**) Temporal image series illustrating the tissue becomes transparent and returns opaque again after experiment. (**c**) Long terms monitoring of the tissue of the mice illustrating the recovery of the mice belly. (**d-e**) Optical imaging results from two different animals. (**f**) Schematics illustrating the application of absorbing molecules on the mouse head. (**g**) Images showing the change of transparency of tissue for visualizing the vessels on the surface of brain. All Scale bars: 5 mm.

**Figure 4.**
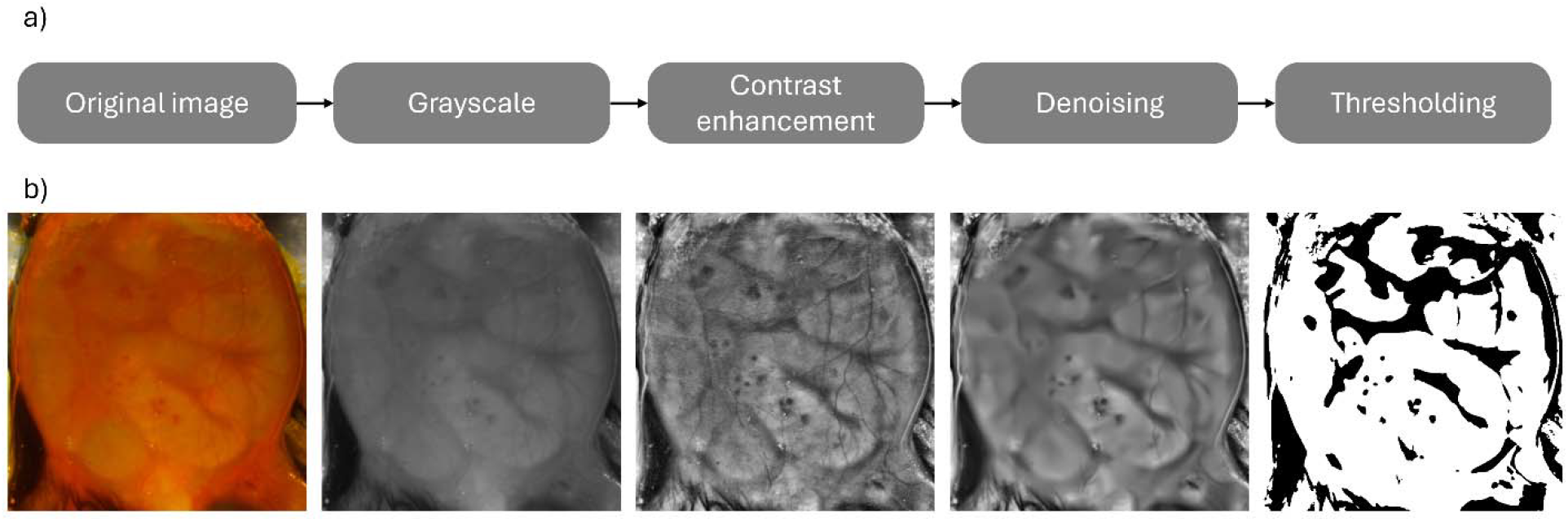
Image processing for improved visualization of organs. (**a**) Workflow showing the image processing steps from a raw image to segmented organ images. (**b**) Representative images showing each step after image processing.

### 4. Increase organ visibility with image processing

#### a) Image processing steps

1. Turning the image to grayscale. The original color image file was loaded and then converted to a grayscale for easier processing of the image. Normalization was performed on the image due to the high-bit-depth of the image (.tif file) and the image was cropped to analyze just the region of interest.
2. Enhance contrast of the image. An intensity distribution analysis was then performed on the image to help understand the pixel intensity spread, detect contrast issues, and identify regions of interest. This was done by plotting a histogram of pixel intensities. To enhance organ visibility, Contrast Limited Adaptive Histogram Equalization (CLAHE) was applied to the image with a clipLimit of 5.0 and a tileGridSize of (16,16) to enable high local contrast. These values were determined through an iterative process until an improved pixel distribution in the histogram was achieved.
3. Denoise the contrast-enhanced image. CLAHE improves contrast but can also lead to an increase in noise in some uniform regions. This can lead to small fluctuations in pixel intensity being misclassified as foreground during segmentation. Non-local means denoising was used for denoising with a filter strength of 11, a patch size of 31 and a window size of 31. Otsu’s adaptive thresholding was applied to the different denoised images.
4. Segment organs of interests based on thresholding. Otsu’s method was chosen because it analyzes the histogram and finds the optimal threshold. This removes the need for trial and error and automatically finds a good cutoff between the organ and the background. To further improve thresholding, a bias of −15 was added to Otsu’s threshold. This causes the algorithm to include low-intensity regions like those beneath the veins.

## REPRESENTATIVE RESULTS

### Demonstration of Optical Transparency in ex vivo chicken breast tissue

To investigate light propagation through ex vivo tissue, we used sliced pieces of raw boneless chicken breast with thicknesses of 1.5 mm, 2.5 mm, and 3.5mm (**Figure 2b**). The tissues were cut parallel to collagen fibers, ensuring consistent muscle fiber orientation. Images were taken using both a mesh grid pattern and “UT Dallas” logo underneath the tissue before and after treating with different solution concentrations (**Figure 2b**). To capture the images, we used a CMOS camera paired with a lens featuring 3.5 mm focal length and f/1.4 aperture, and a LED light source was used for back illumination of the sample. The corresponding transmission data for each tissue was then measured using UV-VIS-NIR spectrometer (**Figure 2c**).

The optical transparency of the tissue can be visualized by the naked eye and quantified with spectrometer measurements. For the untreated chicken tissues, the underlying patterns are completely invisible and the transmission values in the visible region are approximately 2.4%, 1.7% and 1% for tissue thicknesses of 1.5 mm, 2.5 mm and 3.5 mm, respectively. At 1064 nm wavelength, the transmission values are 8%, 5% and 1.8% for same thicknesses. The solutions were prepared by dissolving the Tartrazine powder in deionized water. After the tissues were soaked in Tartrazine solutions of increasing concentrations, the dye molecules diffuse throughout the extracellular matrix and cellular compartments, altering the refractive index of the medium.^9^ This refractive index matching substantially reduces the light scatting within tissue, resulting in enhanced dynamic imaging and higher transmission in spectroscopic measurements. As the Tartrazine concentration increased from 0 to 0.65 M, the refractive index of dye solutions increased, and transparency of the tissue substantially improved. For 0.65 M dye solution, the grid pattern and UT Dallas logo underneath the tissues become clearly visible, and transmission increased to 38.7%, 25%, and 21.7% in the visible spectrum for thicknesses of 1.5 mm, 2.5 mm and 3.5 mm, respectively. At 1064 nm wavelength, the transmission values were 50%, 36.8% and 37.5% for the same samples.

### Demonstration of Optical Transparency in Living animals

For dynamic imaging in live animals, we prepared dye solution with 0.65M concentration and added 0.4% low melting agarose to adjust solution’s viscosity and improve adherence to the skin. During the gut imaging, the animal was anesthetized using inhalation anesthesia and the body temperature was maintained at 37^0^C using a digitally controlled heating pad. Before topical application of 0.65 M dye solution, abdominal fur was removed by shaving and applying hair removal cream on the skin following the process shown in **Figure 3b**. The Tartrazine solution was applied on the skin for 2 minutes and then gently massaged. Immediately, within a few minutes the dye molecules diffuse into the extracellular matrix and superficial dermal layers. The diffusion of the dye molecules reduces refractive index mismatch between low index aqueous interstitial fluid and high index lipid rich structures, resulting in reduced scattering of incident light at tissue interfaces and cellular boundaries. As a result, the skin becomes visibly transparent in visible spectrum for wavelengths longer than 550 nm, making it possible to visualize the deep-seated organs without surgically removing the skin (**Figure 3b-e**). For optical imaging of the brain vessels, the mouse was anesthetized using injectable anesthesia, and fur was removed before the application of dye solution. Our results demonstrate transparency of the scalp and visualization of cerebral blood vessels validating the effectiveness of this technique for brain imaging (**Figure 3f-g**).

To demonstrate the reproducibility of our protocol, we present transparency results obtained for three different mice in **Figure 3**. Young adult female mice (3-4 weeks old, approximately 10-12 g) are used for abdominal imaging including two different mouse lines, C57BL/6J (**Figure 3c,f**) and Chat-Cre x tdTomato (**Figure 3e**). The results obtained for all three mice show enhanced transparency of the skin and visualization of the internal organs after treatment with dye solution, validating this tissue clearing approach. Results from mouse-1 are displayed in detail over multiple time points and reveal gradual transparency over the time of gel solution application while highlighting temporal progression and effectiveness of the method (**Figure 3b-c**). It is worth noting that even for mice with same sex and similar ages, the arrangements and visual appearance of internal organs can vary between individuals, highlighting the unique advantage of chronic imaging of the same animal for developmental studies.

### Reversibility

The optical tissue clearing approach used in this protocol is reversible, which gives this approach a significant advantage for in-vivo applications. The optical transparency of the tissue can be reversed by simply rinsing and gently massaging the skin with water. The image shown in **Figure 3b**, captured after two hours of the abdominal experiment, indicates that transparency is temporary, and it can be fully reversed. Following reversal, the skin maintains normal appearance and mice exhibiting typical activity and hairs on the treated skin regrow within several days (**Figure 3c**). This further confirms the safety and effectiveness of this approach for dynamic imaging in live animals, especially for potential chronic imaging applications of the same animals.

## TABLE OF MATERIALS

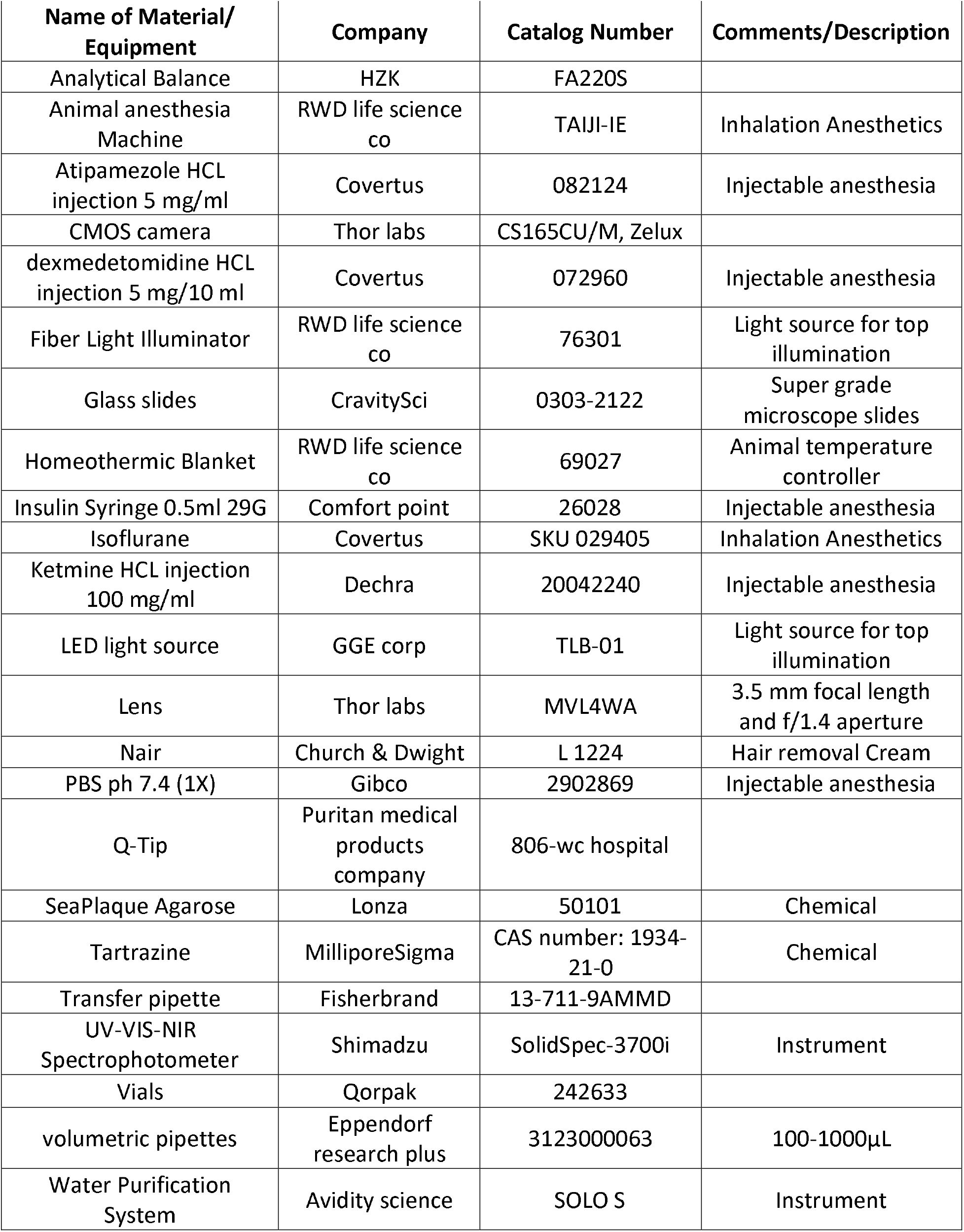

## DISCUSSION

This work presents a comprehensive and reproducible protocol method to achieve optical transparency in biological tissues utilizing absorbing molecules, specifically FDA approved food dye Tartrazine. This method effectively minimizes intrinsic scattering in tissue by altering the refractive index within the tissue’s microenvironment, which enhances optical transparency. We demonstrated the effectiveness of this method through both ex-vivo and in-vivo imaging. For ex-vivo imaging, we investigated the effect of different concentrations of Tartrazine on chicken breast tissue with different thicknesses of the biological samples. Our results indicate that an optimal concentration of Tartrazine can make chicken breast tissues transparent to the naked eye. The spectroscopic measurements of transmission further reveal that although incident is almost completely absorbed below 550 nm, transmission of light through the tissue significantly increases for longer wavelengths of visible spectrum and into the near infrared region. Our current protocol relies on the passive diffusion of dye molecules into the tissue, which limits the applications to larger samples, since the diffusion time required is proportional to the square of the thickness of the samples.^2^

For in-vivo imaging, we have demonstrated this approach for abdominal and brain imaging in live animals. Our innovative transient optical clearing technique has led to a new strategy for integrative biology research by enabling high-resolution, real-time imaging of dynamic anatomical and functional processes, such as organ regeneration and internal movement, within intact live tissues. This method is compatible with existing optical imaging modalities and contrast agents, noninvasive, and allows imaging at any location within the animal body without restricting animal behavior. Combined with other optical imaging techniques such as multiphoton-microscopy^18^ and near-infrared imaging,^19^ it can provide detailed insights into structural characterization, microenvironment, and cell phenotypes, facilitating studies on dynamic physiological processes and disease prevention.

## ACKNOWLEDGMENTS

ZO acknowledges support from The University of Texas at Dallas.

## REFERENCES

1. Yun, S.H. Kwok, S.J.J. Light in diagnosis, therapy and surgery. Nat. Biomed. Eng. 1 (1), 0008 (2017).

2. Tuchin, V. V. Optical clearing of tissues and blood. (SPIE Publications, 2005).

3. Ertürk, A. Deep 3d histology powered by tissue clearing, omics and ai. Nat. Methods. 21 (7), 1153–1165 (2024).

4. Chung, K. et al. Structural and molecular interrogation of intact biological systems. Nature. 497 (7449), 332–337 (2013).

5. Costantini, I., Cicchi, R., Silvestri, L., Vanzi, F., Pavone, F. S. In-vivo and ex-vivo optical clearing methods for biological tissues: Review. Biomed. Opt. Express. 10 (10), 5251– 5267 (2019).

6. Genina, E. A. Tissue optical clearing: State of the art and prospects. Diagnostics. 12 (7), 1534 (2022).

7. Ueda, H. R. et al. Tissue clearing and its applications in neuroscience. Nat. Rev. Neurosci. 21 (2), 61–79 (2020).

8. Mai, H. et al. Whole-body cellular mapping in mouse using standard igg antibodies. Nat. Biotechnol. 10.1038/s41587-023-01846-0 (42), 617–627 (2024).

9. Ou, Z. et al. Achieving optical transparency in live animals with absorbing molecules. Science. 385 (6713), eadm6869 (2024).

10. Sai, T., Saba, M., Dufresne, E. R., Steiner, U., Wilts, B. D. Designing refractive index fluids using the kramers–kronig relations. Faraday Discuss. 223 (0), 136–144 (2020).

11. Yu, T. Zhu, D. Strongly absorbing molecules make tissue transparent: A new insight for understanding tissue optical clearing. Light Sci. Appl. 14 (1), 10 (2025).

12. Drexler, W. et al. in Optical coherence tomography: Technology and applications 10.1007/978-3-319-06419-2_10 eds Drexler, W. Fujimoto, J.G.) 277–318, Springer International Publishing, Cham (2015).

13. Miller, D. A. et al. Enhanced penetration depth in optical coherence tomography and photoacoustic microscopy in vivo enabled by absorbing dye molecules. Optica. 12 (1), 24–30 (2025).

14. Zuo, T., Tao, C., Liu, X. Absorbing molecules as optical clearing agents improve the resolution and sensitivity of photoacoustic microscopy. Opt. Lett. 50 (7), 2282–2285 (2025).

15. Wang, F. Zhou, C. ‘Transparent mice’: Deep-tissue live imaging using food dyes. Commun. Bio. 7 (1), 1307 (2024).

16. Yu, H., Liu, J.-B., Ma, Y.-S., Fu, D. Transparent live mice offer more possibilities for biomedical research. Innov. Med. 10.59717/j.xinn-med.2024.100098100098 (2024).

17. Shabbir, M. W., Biswas, S., Kajla, R., Nadella, S., Ou, Z. Designing refractive index fluids of food dye for light propagation through scattering media. arXiv preprint 2503.05105. (2025).

18. Scheele, C. L. G. J. et al. Multiphoton intravital microscopy of rodents. Nat. Rev. Methods Primers. 2 (1), 89 (2022).

19. Schmidt, E. L. et al. Near-infrared ii fluorescence imaging. Nat. Rev. Methods Primers. 4 (1), 23 (2024).

